# Low *XIST* expression in Sertoli cells of Klinefelter syndrome patients caused the high susceptibility of these cells to an extra X chromosome

**DOI:** 10.1101/2020.08.26.269423

**Authors:** LiangYu Zhao, Peng Li, ChenCheng Yao, XiaoYu Xing, Chao Yang, JiaQiang Luo, ZhiYong Ji, RuHui Tian, HuiXing Chen, ZiJue Zhu, ZhiWen Deng, Na Li, Jing Fang, Jie Sun, ChenChen Wang, Zhi Zhou, Zheng Li

## Abstract

Klinefelter syndrome (KS) is the most common genetic cause of human male infertility. Patients suffer from heterogeneous testicular atrophy with loss of both germ cells and Sertoli cells. However, the mechanism by which the extra X chromosome causes failure of spermatogenesis remains poorly understood. Here, we profiled testicular single-cell transcriptomes from three KS patients and compared the results with those of healthy donors. Among different somatic cells, Sertoli cells showed the greatest changes in KS patients. Further analysis showed that *XIST*, a key long intergenic non-coding RNA that inactivates one X chromosome in female mammals, was widely expressed in somatic cells, except for Sertoli cells, leading to an increase in X-inactivation genes in these cells, which may cause Sertoli cells death and disruption of the spermatogenic microenvironment. Our study proposed a new mechanism to explain the unique pathological manifestations of KS in the testes and provided a theoretical basis for subsequent research and related treatment.

## Introduction

Klinefelter syndrome (KS), 47, XXY, is one of the most frequent genetic causes of infertility, occurring in approximately 0.15‰ of newborn males(Bojesen et al. 2003). KS patients exhibit persistent and irreversible degeneration of seminiferous tubules in the testes, which is manifested as a significantly reduced testicular volume since puberty, and the difference becomes even greater by adulthood (Aksglaede et al. 2011). Within the testes of KS patients, Sertoli cells, as the scaffold of the seminiferous tubules, are completely absent in most areas. However, the remaining Sertoli cells can still construct a seminiferous tubule structure (although the structure exhibits obvious pathological morphology, including reduced cell volume, an increased nucleo-cytoplasmic ratio, and loss of intercellular space) with other somatic cells and even support complete spermatogenesis(Bergère et al. 2002), indicating that the dose effect of the X chromosome genes greatly affects the cell fate and testicular morphology. Furthermore, compared with Sertoli cells, the damage caused by the extra X chromosome in other testicular cells (such as Leydig cells) and other systems (such as the cardiovascular, hepatobiliary, and respiratory system) is so mild in KS patients that it is often neglected by doctors, indicating that different types of cells also show susceptibility for X chromosome overload.

The activation and inactivation of the X chromosome (XCI) are regulated by completely different mechanisms in various biological processes. From embryonic day 7.5 (E 7.5) to E 12.5 of development, primordial germ cells undergo global removal of DNA methylation, leading to a global (including the X chromosome) increase in transcription(Popp et al. 2010). During meiosis, the unsynapsed X and Y chromosomes are transcriptionally silenced through meiotic sex chromosome inactivation and are sequestered in a cellular domain known as the sex body(Handel 2004). After initiation of spermiogenesis, the histone-to-protamine transition and high level of DNA methylation in spermatids also lead to a global or an independent repression of sex chromosome expression(Namekawa et al. 2006; Sin et al. 2012). In addition to the above mechanisms of X chromosome silencing during spermatogenesis, *XIST*-dependent XCI, a more universal XCI mechanism, plays a critical role in balancing the gene dosage between XY and XX cells via silencing of one of the two X chromosomes in XX cells(Carrel and Willard 2005; Galupa and Heard 2015).

However, it is certain that there always are some genes are not or are only slightly affected by XCI process(Mueller et al. 2013; Sin and Namekawa 2013). These genes escape X inactivation: in women about 15% of X-linked genes are bi-allelically expressed and most genes have a stable inactivation pattern(Berletch et al. 2011). Expression from the inactive X allele varies from a few percent of that from the active allele to near equal expression(Naumova et al. 2013; Robert Finestra and Gribnau 2017). Therefore, we hypothesized that heterogeneity of X chromosome overload damage in different KS cells may occur because of the failure of the XCI process (a global increase in X gene expression) or the overexpression of some X genes that escape inactivation. Therefore, we profiled and analyzed over 37,000 single cell transcriptomes from whole testes of patients with obstructive azoospermia (OA, patients with normal spermatogenesis) and KS patients and compared the X gene expression pattern among different testicular cells and within the same cell type. In addition, we also attempted to explain the etiological characteristics of several known XCI mechanisms to increase the understanding of the pathogenesis of azoospermia caused by KS.

## Results

### Global transcriptome profiling showed that Sertoli cells underwent the greatest change in KS patients

To understand the global changes in each type of testicular cell in KS patients, we merged the single cell expression data of the KS (3 patients) group with those of the OA (2 patients) and YqAZFa microdeletion (AZFa_Del, 1 patient) groups. The sex hormone levels of two OA donors were within the normal range. All three KS and donors and the AZFa_Del donor had higher levels of follicle-stimulating hormone (FSH) and luteinizing hormone (LH), but their testosterone (T) and estradiol levels were in the normal range (Figure S1A). In total, 27,597 OA cells, 8267 KS cells and 5434 AZFa_Del cells were profiled. On average, 9359 reads (unique molecular identifiers, UMIs), 2719 expressed genes, and 3.7% mitochondrial genes were detected per cell; however, different types of cells showed high heterogeneity. In general, germ cells exhibited more active transcription and a lower proportion of mitochondrial genes (Figure S2A). T-distributed stochastic neighbor embedding (tSNE) analysis showed that cells from the three types of patients had obviously different 2D spatial distributions, but good repeatability was demonstrated within the patient groups (Figures 1B and S1C). In total, nine cell clusters were identified in the whole cell population based on the expression of known cell type-specific markers (Figure 1C and S2B).

**Figure 1.**
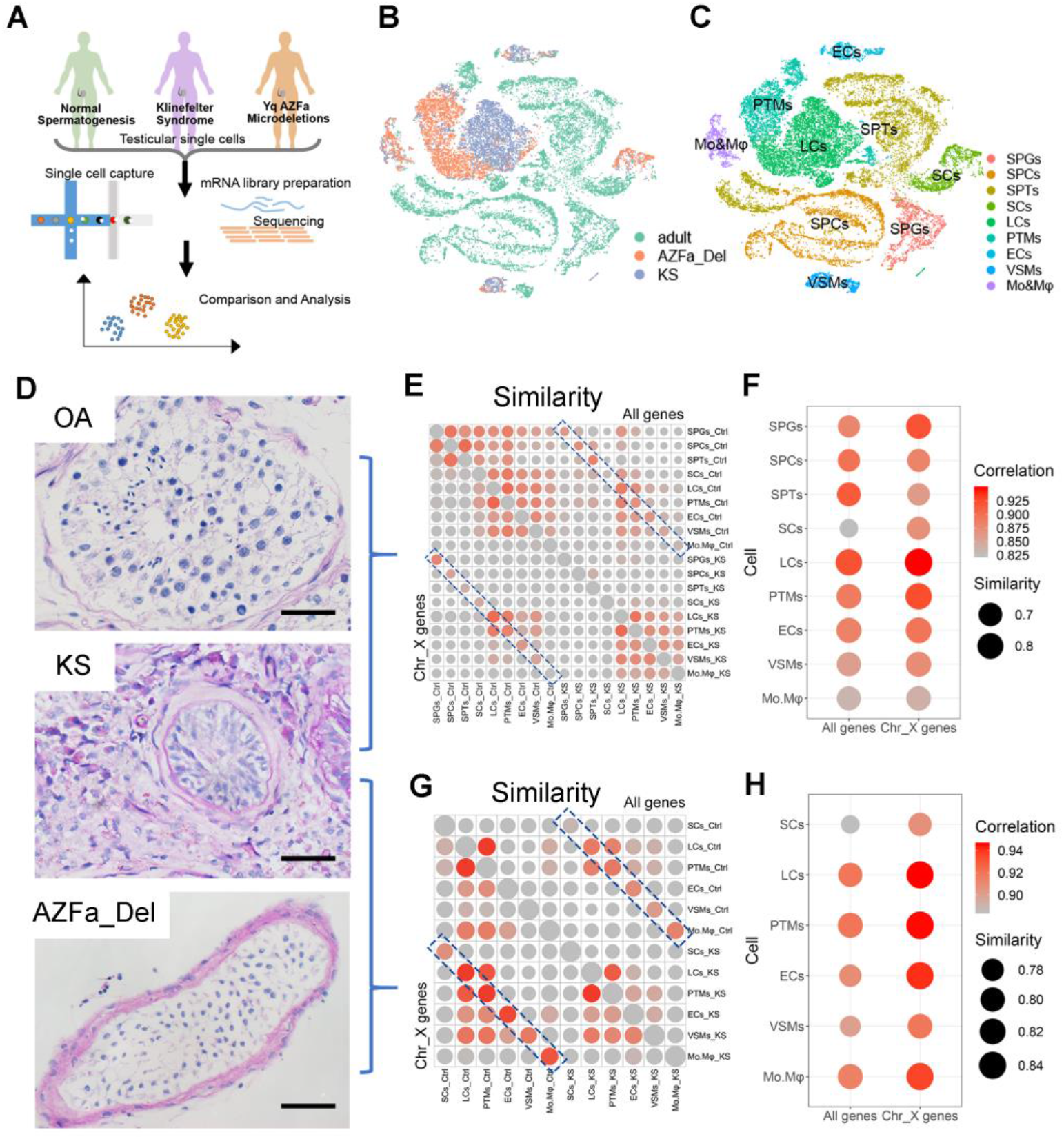
The global transcriptome pattern of obstructive azoospermia, Klinefelter syndrome, and AZFa_Del testes. (A) Schematic illustration of the study workflow. (B–C) T-distributed stochastic neighbor embedding plots of all six testicular single-cell samples. Each cell is labeled with a different color according to its sample type (B) or cell cluster (C). (D) Glycogen periodic acid Schiff staining of obstructive azoospermia (OA), Klinefelter syndrome (KS), and AZFa_Del testes illustrated typical pathological changes in their histological morphology. The scale bar represents 50 μm. (E–H) A bubble diagram shows the similarity between KS and OA (E) or AZFa_Del cells (G). The upper right half represents the result when all expressed genes are considered, and the lower left half represents the results when only X chromosome genes are considered. The right panel shows an enlarged view of the contents of the blue dotted rectangle, illustrating the similarities between the same types of KS and OA (F) or AZFa_Del (H) cells. The size of points represent the value of Jaccard distance. The colors of the points represent the regression coefficients.

The periodic acid Schiff (PAS)/hematoxylin staining results showed that the OA testes exhibited normal morphology. In the AZFa_Del testis, seminiferous tubules showed a Sertoli cell-only syndrome with a Johnsen score of 2 and exhibited vascular basement membrane thickening without interstitial proliferation. Additionally, the morphology of these Sertoli cells was similar to that of OA testes. Unlike other types of nonobstructive azoospermia, the typical pathological changes in different regions of KS testes showed significant heterogeneity. In KS testes, most seminiferous tubules exhibited more severe atrophy, with Johnsen scores of 0–1 (germ cells were completely absent, and Sertoli cells were also reduced with abnormal morphology) and exhibited vascular basement membrane thickening with severe interstitial proliferation; however, a small number of larger and opaque seminiferous tubules with incomplete or even normal spermatogenesis were still present (Figure 1D and S1B).

Then, to further assess the effects of an extra X chromosome on various cell types, we calculated the Jaccard distance and correlation among nine cell types between KS and OA patients and found that Sertoli cells had the least similarity (largest Jaccard distance) and lowest correlation (smallest r index) compared with other cell types (Figure 1E and 1F). When only the X chromosome genes were considered, Sertoli cells were still one of the most variable cell clusters (Figure 1F). In addition, to exclude the potential effects of germ cells on somatic cells, we also compared KS cells with AZFa_Del cells using the same method and obtained a similar result (Figure 1G and 1H). These results indicated that Sertoli cells underwent the greatest change of any cell type in the presence of an extra X chromosome.

### Transcriptional changes in KS Sertoli cells mainly occurred on the X chromosome genes

Because Sertoli cells underwent the greatest change among nine cells clusters in KS patients, we then focused on these cells and further analyzed the changes that occurred. The global transcriptional level of Sertoli cells was lowest among other cell types. The total number of expressed genes and total UMI count per cell in KS Sertoli cells were higher than those in OA Sertoli cells. This result was also observed in germ cells but not in other somatic cells (Figure 2A and 2B). When the fold change threshold was set to 0.25, we observed 365 downregulated and 1041 upregulated differentially expressed genes (DEGs) in KS Sertoli cells compared with OA Sertoli cells (Table S1), and most of these DEGs were located on chromosome 1 (139 genes) and 19 (115 genes) and the X chromosome (87 genes) (Figure 2C). When the fold change threshold was increased to 0.5, most of DEGs were still located on these chromosomes. However, when the fold change threshold was further increased to 1, only 29 DEGs were found, and seven were located on the X chromosome; this proportion was much higher than that observed for autosomes (Figure 2D). This result indicated that the extra X chromosome had a significant dose effect on gene expression levels; however, the dosage effect in other somatic cells was obviously weaker than that in Sertoli cell (Figure 2E). As expected, these seven X chromosome DEGs were increased in KS patients compared with OA patients. Interestingly, among these X chromosome DEGs, *CD99* and *SLC25A6* escaped from XCI (eXi genes); however, *RPL36A, TMSB4X, RPL39*, and *BEX1* did not escape from XCI (neXi genes) according to a previous study on females (Figure 2F). These results indicated that some neXi genes also escaped from XCI in KS Sertoli cells, leading to an increased X chromosome expression level in KS Sertoli cells.

**Figure 2.**
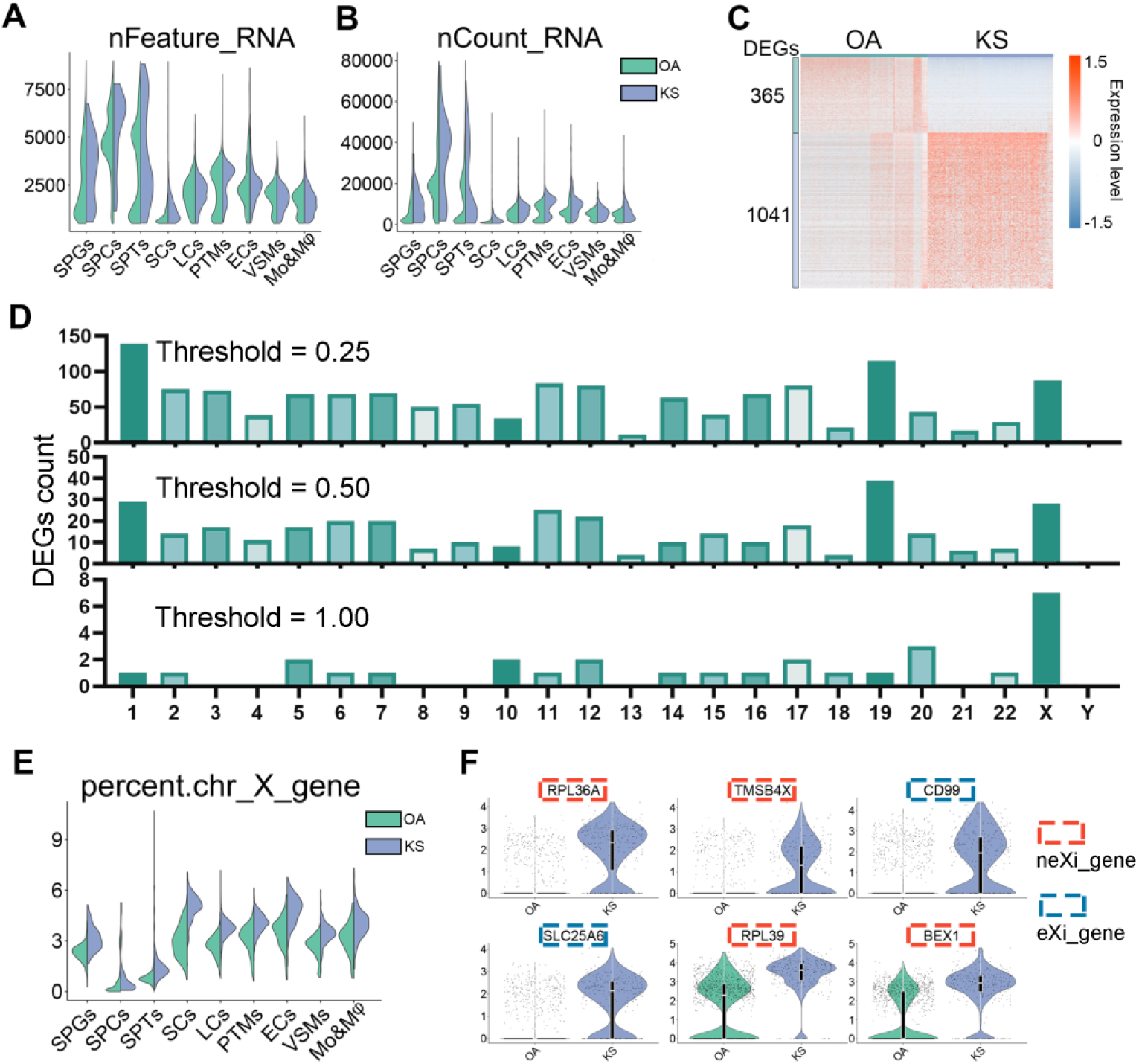
Comparison of global and X chromosome gene expression patterns between obstructive azoospermia and Klinefelter syndrome Sertoli cells. (A–B) Violin plot shows the total expression gene counts and unique molecular identifier counts of Sertoli cells; the column is divided according to cell type, including obstructive azoospermia (OA) or Klinefelter syndrome (KS) cells. (C) Heatmap shows differentially expressed genes (DEGs) between OA and KS Sertoli cells. DEG counts are shown on the left of the color bar, which indicates the cell type annotation. (D) The number of DEGs between OA and KS Sertoli cells located on each chromosome under different thresholds. (E) A violin plot shows the percentage of X chromosome gene expression in Sertoli cells; the column is divided into OA and KS cells. (F) Violin plot shows the expression of the top six DEGs located on the X chromosome of Sertoli cells. The color of the rectangular box represents whether this gene can escape from X chromosome inactivation according to a previous study on females.

### Failure of XCI in Sertoli cells

To further understand why KS Sertoli cells underwent greater increase in the X gene level compared with OA Sertoli cells among somatic cells, we calculated the percent of eXi gene and neXi gene expression in nine clusters of cells. As expected, the percentage of eXi genes in all six types of KS somatic cells increased at least 1.5-fold but less than 2-fold compared with OA cells, indicating that the X chromosome gene expression level and X chromosome number were not strictly linearly correlated. However, the percentage of neXi gene expression in KS Sertoli cells was also 1.5 times higher than that in OA cells, and this result was not found in the other five somatic cell types (even though the percentage of neXi gene expression in these cells was also increased in KS cells). All of these results indicated that Sertoli cells have different levels of ability to silence the extra X chromosome compared with other somatic cells (Figure 3A, 3B, and 3C).

**Figure 3.**
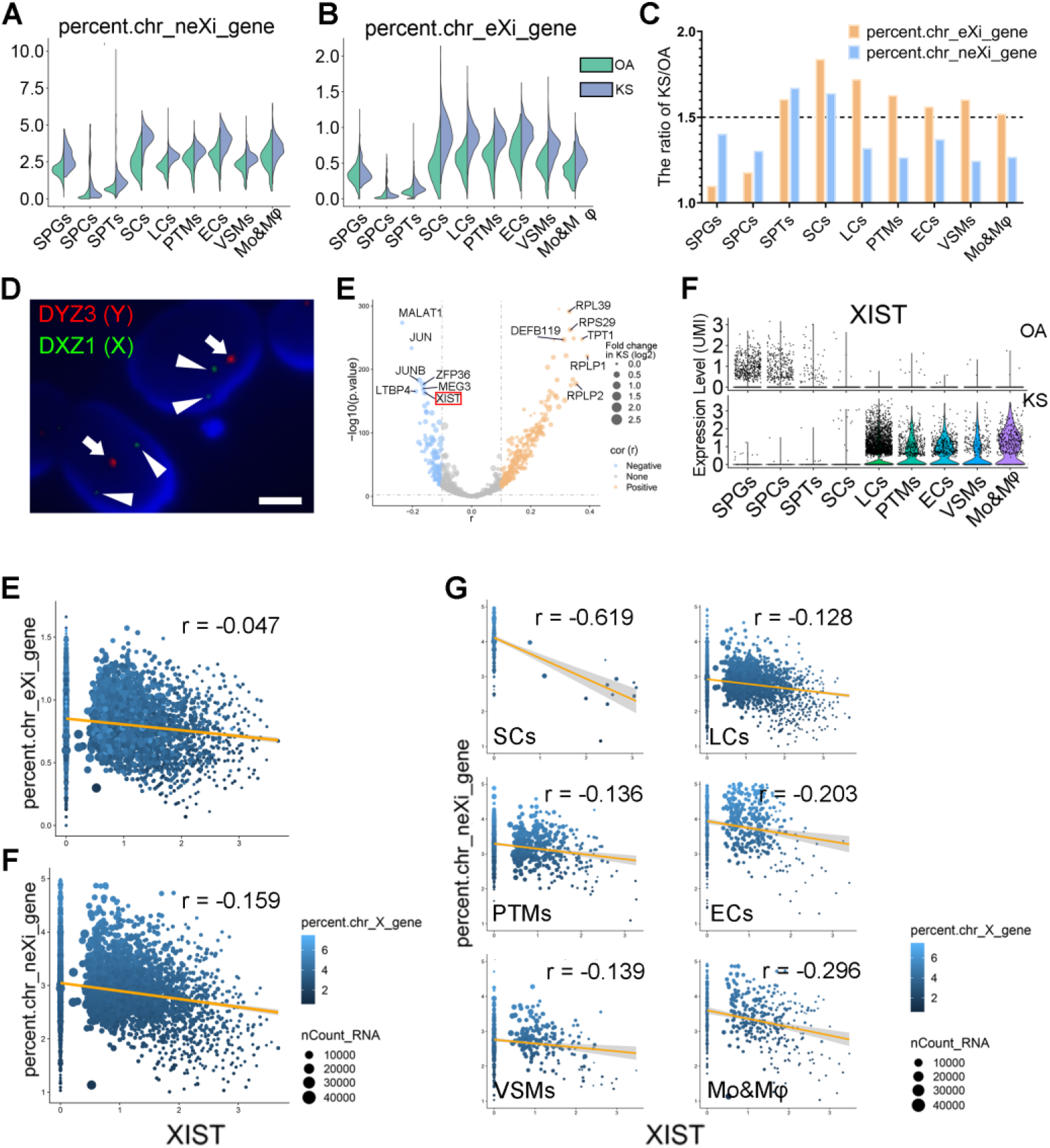
The relationship between X chromosome gene expression patterns and *XIST* expression levels in Klinefelter syndrome. (A–B) Violin plot shows the percent of genes that did not escape from X chromosome inactivation (XCI) (neXi genes) (A) and those that escaped from XCI (eXi genes) (B) in Sertoli cells. The column is divided according to cell type, including obstructive azoospermia (OA) or Klinefelter syndrome (KS). (C) The ratio of neXi gene or eXi gene expression levels between KS and OA cells. (D) DNA-fluorescence in situ hybridization shows the karyotype of sex chromosomes of KS Sertoli cells. The long arrow indicates the DYZ3 region of the Y chromosome (labeled by green fluorescence), and the triangular arrow indicates the DXZ3 region of the X chromosome (labeled by red fluorescence). The scale bar represents 0.5 μm. (E) Volcano plot shows the correlation between neXi gene expression and that of each gene in all KS testicular cells. The X-axis represents the r value of the correlation coefficient. The Y-axis represents the P value. The sizes of the points represent the fold change in the gene expression level in KS cells compared with OA cells. (F) Violin plot shows the expression level of *XIST* in each cell cluster of OA (upper panel) or KS (lower panel) cells. (E–G) Regression analysis shows the correlation between *XIST* expression and the eXi (E) or neXi (F) gene expression level in total KS somatic cells or in different clusters of KS somatic cells (G). The sizes of the points represent the total unique molecular identifier count of the cells, and the color represents the percentage of X chromosome gene expression.

Next, we identified the karyotype of KS Sertoli cells to determine whether the reason for this phenomenon was that KS Sertoli cells contained more than two X chromosomes. However, the DNA-fluorescence in situ hybridization (FISH) results showed that the karyotype of all Sertoli cells of all three patients in this study was XXY (Figure 3D), indicating that KS Sertoli cells do lose the XCI function. Next, we calculated the correlation between the expression level of each gene and the percentage of neXi gene expression in KS somatic cells (including Sertoli cells) to identify the universal upstream or downstream regulatory genes that are most closely related to XCI. As the result, *MALAT1, JUN, JUNB, LTBP4, ZFP36, MEG3*, and *XIST* showed the highest negative correlation with neXi gene expression (Figure 3E). *XIST* was the first non-coding gene identified within the X inactivation center (XIC), is expressed exclusively from the XIC of the inactive X chromosome, and is essential for the initiation and spread of XCI. Except for Sertoli cells, KS somatic cells had a significantly higher level of *XIST* than OA cells (Figure 3F). In somatic cells (including Sertoli cells), the *XIST* expression level was negatively correlated with the percentage of neXi gene expression (r=-0.159); however, its correlation with the percentage of eXi genes was not strong (r=-0.047). Although the expression level of *XIST* was low in KS Sertoli cells, to answer the question of whether *XIST* affects neXi gene expression in these cells, we isolated KS Sertoli cells and found that *XIST* expression was strongly negatively correlated with the percentage of neXi gene expression (r=-0.619), and this negative correlation was also observed in other somatic cells (Figure 3G). These results indicate *XIST* is a universal silencing factor for neXi gene expression in KS somatic cells, including Sertoli cells, and the abnormal increase in neXi gene expression in KS Sertoli cells may be caused by low *XIST* expression.

### Identification of known and novel candidate regulators of *XIST* expression in KS testicular cells

To explore why KS Sertoli cells expressed the lowest level of *XIST* among the nine clusters, we further predicted and analyzed the regulators of *XIST*. In total, 20 candidate regulators were found via the PROMO dataset, and *FOXA1, PAX5, NF1, CEBPB*, and *GATA2* showed the top binding scores for the promotor region of *XIST* (Figure 4A). Among these new candidates and other known regulators (such as *MYC, JPX*, and *KLF4), CEBPB, HOXD9, AR*, and *MYC* were upregulated in KS cells, while *TBP*, *NF1*, and *DNMT1* were downregulated in KS cells (Figure 4B). After regulators with low expression levels were filtered out, we found eight new candidate regulators that showed a cell type-specific expression pattern. Specifically, *YY1*, *NR3C1*, and *TCF7L2* were expressed in both germ cells and somatic cells, and *CREBPB, GATA2, HOXD9, IRF2*, and *ETS* were only expressed in one or some types of somatic cells. Interestingly, all of the predicted regulators showed no or extremely low expression in Sertoli cells (Figure 4C). In addition, some reported regulators of *XIST* also showed an “absent in Sertoli” expression pattern (Figure 4D). However, the expression of most of these regulators was higher in somatic cells than in germ cells. The cell type-specific pattern of these *XIST* regulators may be related to the loss of *XIST* expression in Sertoli cells and germ cells.

**Figure 4.**
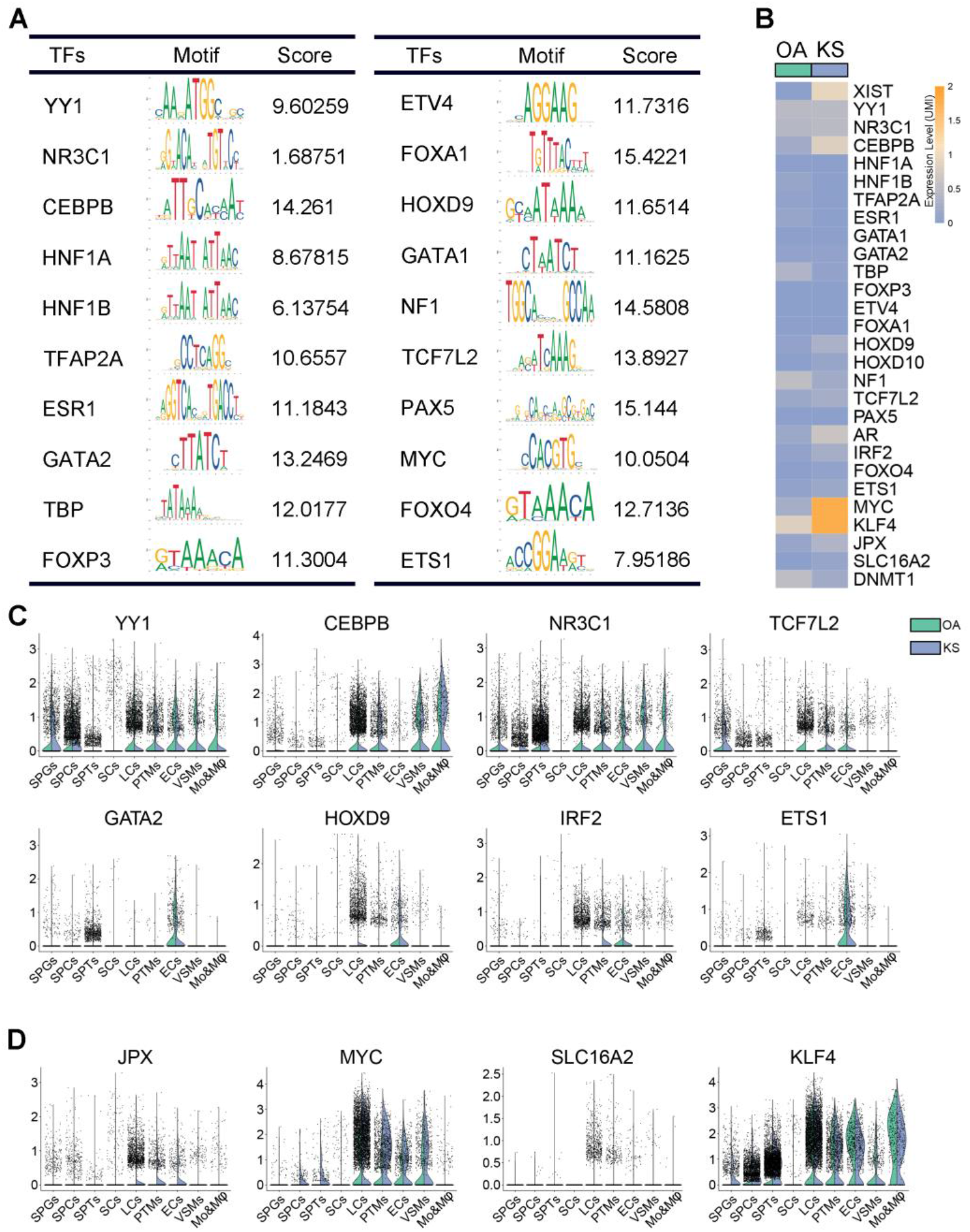
Candidate regulators of *XIST*. (A) Twenty candidate transcription regulators and their binding motifs within −3 kb of the transcriptional start site of the *XIST* gene. The score was calculated using the Jaspar dataset. (B) Heatmap showing the expression level of *XIST* and its regulators in obstructive azoospermia (OA) and Klinefelter syndrome (KS) somatic cells. (C–D) A violin plot shows the expression level of *XIST* regulators predicted by the PROMO dataset (C) or validated by previous studies (D) in each cell cluster of OA and KS cells.

### The pathological changes of biological process in KS Sertoli cells

Sertoli cells function as scaffolds of seminiferous tubules and “nurse” cells of spermatogenesis and therefore are one of the most important parts of the spermatic environment. The pathological changes in KS Sertoli cells were directly related to the phenotype of KS testes (seminiferous tubule atrophy and failure of spermatogenesis). To understand these changes, we first predicted the upstream signaling with ingenuity pathway analysis (IPA) based on the DEGs between OA and KS Sertoli cells. Among these signaling pathways, we found that metribolone (a synthetic non-aromatizable androgen and anabolic steroid), β-estradiol, and interleukin (IL)-5 contributed to a strong activation effect (activation z score >2) in KS Sertoli cells, indicating that an abnormal inflammatory response and hormone disruption occurred in these cells. Consistent with signaling activation in the KS phenotype, immunosuppressant-associated drugs such as sirolimus(Sehgal 2003) and CLPP(Bhandari et al. 2018) produced negative signaling with regard to the KS phenotype in Sertoli cells (Figure 5A). Next, we analyzed the changes in pathways in KS Sertoli cells, and IPA results showed that “estrogen receptor signaling” and inflammatory response terms such “CXCR4 signaling”, “Fcγ receptor-mediated phagocytosis in Mo&Mφ”, and “IL-8 signaling” were activated in KS Sertoli cells (Figure 5B). Considering that some KS patients have symptoms of androgen deficiency (the androgen levels of all three KS patients were in the low to normal range, but the gonadotrophic hormone levels were higher than the normal range), abnormal enrichment of “estrogen receptor signaling” in KS Sertoli cells may also be caused by Leydig cell dysfunction. Therefore, we also examined changes in the androgen synthesis pathway in KS Leydig cells. Interestingly, most genes related to testosterone and dihydrotestosterone synthesis were upregulated in KS cells compared with normal adult cells (Figure S3A), which could be caused by the high gonadotropin levels in KS patients. This result indicated that low androgen levels in KS patients were caused by a decreased number of Leydig cells rather than dysfunction of androgen synthesis in these cells.

**Figure 5.**
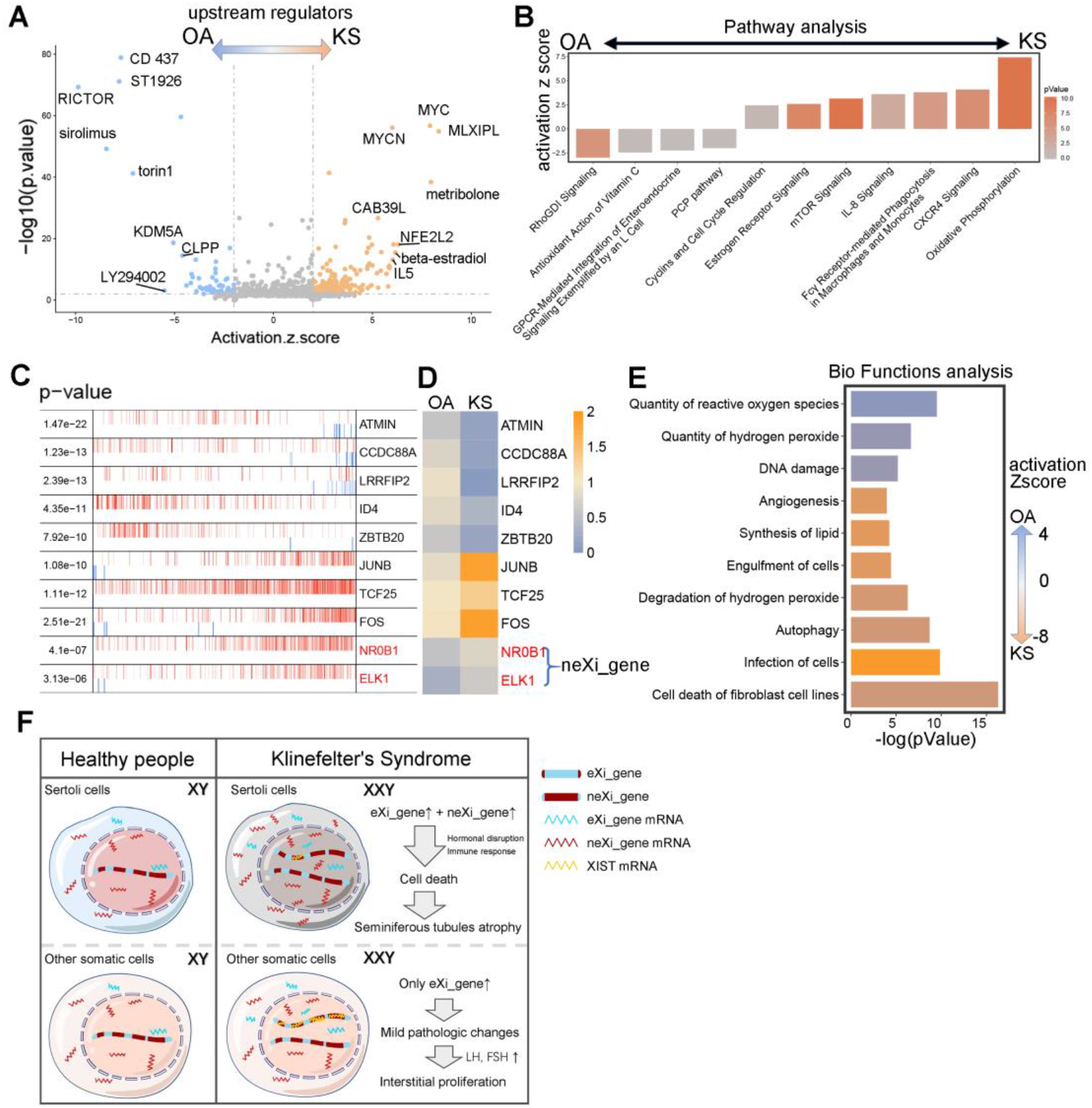
Changes in the regulatory pathway and bio-function in Klinefelter syndrome Sertoli cells. (A) Volcano plot shows upstream signaling, which may induce the differences between obstructive azoospermia (OA) and Klinefelter syndrome (KS) Sertoli cells. (B) Ingenuity pathway analysis (IPA) pathway terms that are enriched and decreased in KS Sertoli cells are shown as a bar plot. The Y-axis represents the z score, and the red gradient indicates low to high P values. (C–D) The top 10 candidate master regulators that may induce the differences between OA and KS Sertoli cells were identified by the master regulator analysis algorithm (MARINa). (C) Heatmap showing the expression levels of those regulators in OA and KS Sertoli cells. In the MARINa plots, activated targets are colored red, and repressed targets are colored blue for each potential master regulator (vertical lines on the X-axis). On the X-axis, genes were rank-sorted by their differential expression in OA and KS Sertoli cells from left to right. The P values on the left indicate the significance of enrichment. (E) IPA bio-function terms enriched or decreased in KS Sertoli cells are shown as a bar plot. The X-axis represents the P value, and the gradient from orange to blue indicates KS to OA bias activation. (F) The schematic diagram shows the main pathologic changes and associated mechanism in KS somatic cells.

To investigate the master regulators and construct the transcriptional regulatory network during the pathological state of KS, we utilized the Algorithm for the Reconstruction of Accurate Cellular Networks (ARACNe) method to analyze all 1469 known transcription factors from the Animal Transcription Factor Database. Compared with OA Sertoli cells, *ATMIN, CCDC88A, LRRF/P2, ID4*, and *ZBTB20* expression levels were decreased, and *JUNB, TCF25, FOS, NR0B1*, and *ELK1* expression levels were increased in KS Sertoli cells (Figure 5C and 5D), indicating possible regulatory roles in the pathological process.

When specific bio-function changes were examined, we noticed that terms such as “cell death of fibroblast cell lines”, “infection of cells”, “autophagy”, and “synthesis of lipid” were enriched in KS Sertoli cells, and terms such as “quantity of reactive oxygen species”, “quantity of hydrogen peroxide”, and “DNA damage” were enriched in OA Sertoli cells (Figure 5E). In general, these results described the changes in upstream signaling, pathways, regulators, and bio-functions in KS Sertoli cells (Figure 5F).

### KS germ cells with low *XIST* expression were in the maintenance phase of XCI

In the above analysis, we noticed that both Sertoli cells and germ cells expressed low *XIST* levels (Figure 3F); however, the percentage of X chromosome genes in spermatogonia (SPGs) and spermatocytes (SPCs) in KS patients were not as high as those in KS Sertoli cells. To explain this paradox, we further analyzed the other mechanisms in germ cells that may affect X chromosome activation. Germ cells were further divided into 11 clusters according known markers (Figure 6A and 6C), and each cluster contained KS cells, which showed little difference from OA cells (Figure 6B). Interestingly, *XIST* was mainly expressed in spermatogonia stem cells (SSCs), and the *XIST* expression level in KS cells was not higher than that in OA cells (Figure 6D). Unlike KS somatic cells, no significant correlation was found between the *XIST* expression level and the percentage of neXi gene expression, indicating that *XIST* may not play a key role in XCI in KS SSCs (Figure 6E). In addition, the change in the expression level of sex chromosome genes during germ cell development was far greater than the difference between KS and OA cells. During meiosis, the expression of sex chromosome genes was extremely limited in both OA and KS cells (Figure 6F and 6G), indicating that the meiotic silencing of unsynapsed chromatin (MSUC) mechanism has an important and effective function in both X chromosomes in KS germ cells. Regarding the SSCs stage, we found no difference in neXi gene expression levels between OA and KS cells. Therefore, we calculated the correlation between the percentage of neXi gene expression and the expression level each gene in KS SSCs and found that *CXorf23, CSAD, ACO2, PTGDS, NOVA1*, and *IGFBP7* showed a negative correlation (Figure 6J and 6K). In addition, the expression levels of *CSAD, PTGDS, NOVA1*, and *IGFBP7* in KS SSCs were significantly higher than those in OA SSCs, indicating that these genes may be involved in the feedback regulation of *XIST* transcription in KS SSCs. In addition to these candidate regulators of *XIST*, we also noticed that JARID2, which is implicated in the initial *XIST*-induced targeting of PRC2 to the inactive X chromosome(da Rocha et al. 2014), showed a significant correlation with X chromosome gene expression levels in KS SSCs (Figure 6J). During the initiation and maintenance phases of XCI, the PRC1 and PRC2 complexes were recruited and bind to the X chromosome and helped to lock in the inactive state of the X chromosome(Sarma et al. 2014). Interestingly, nearly all parts of the PRC1 and PRC2 complexes showed a germ cell-enriched expression pattern, and *SFMBT1, BMU, EZH2*, and *SUZ12* exhibited low expression levels in somatic cells (Figure 6M). In addition, *SLC25A3* and *RBBP4* were highly expressed in KS Sertoli cells but not in normal Sertoli cells (Figure 6M). Based on these results, we speculated that even though *XIST* exhibited low expression levels in both KS Sertoli cells and germ cells, the latter still showed a relatively low level of X gene expression, which may be associated with a germ cell-specific PRC1/2-dependent XCI mechanism.

**Figure 6.**
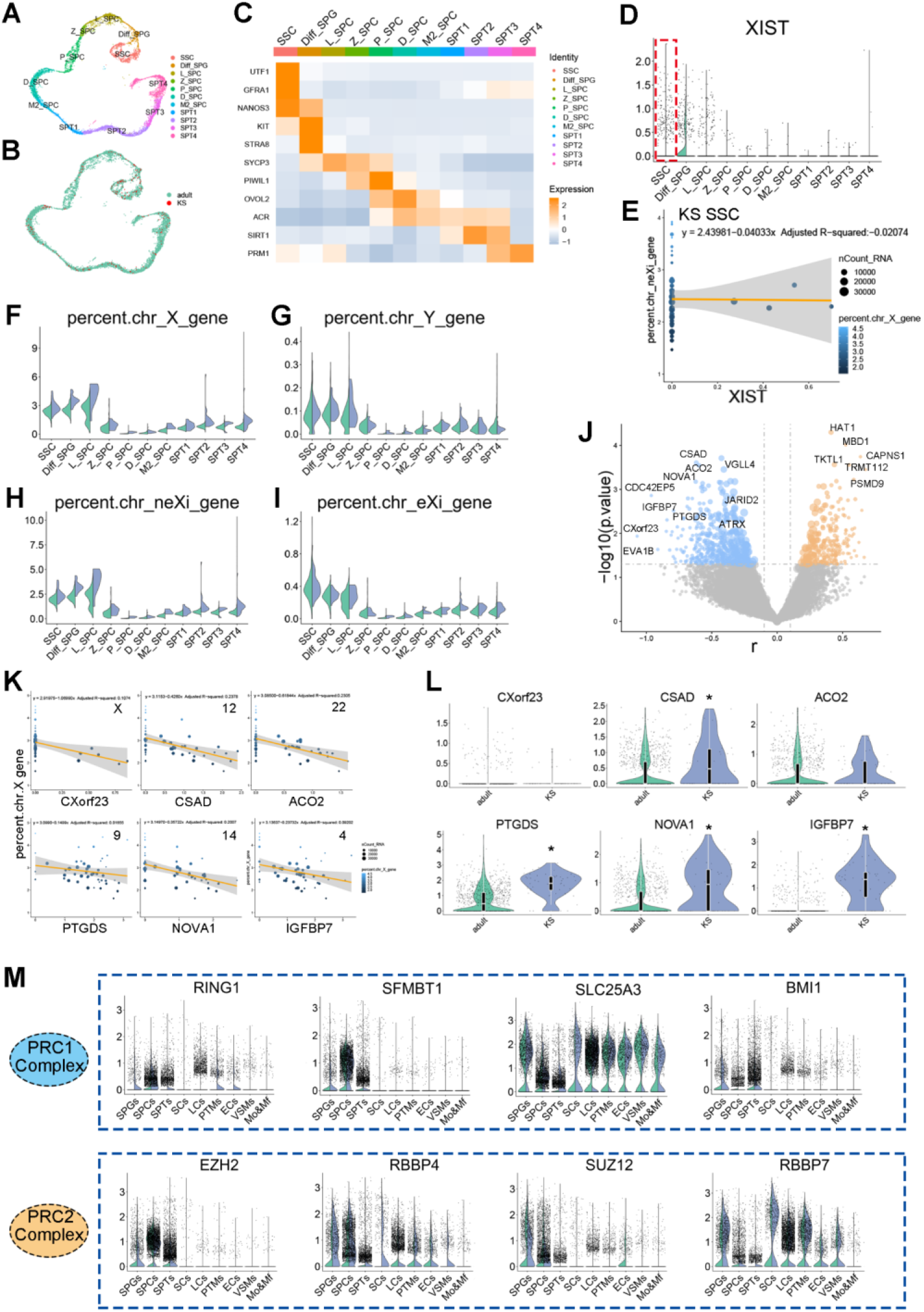
The regulation of X chromosome gene expression patterns in Klinefelter syndrome germ cells. (A–B) T-distributed stochastic neighbor embedding plots of obstructive azoospermia (OA) and Klinefelter syndrome (KS) germ cells. Each cell is labeled with a different color according to its cell cluster (A) or sample type (B). (C) Heatmap showing the expression level of known stage-specific markers during human spermatogenesis in OA and KS germ cells. (D) Violin plot shows the expression level of *XIST* in each germ cell subpopulation of OA and KS cells. (E) Regression analysis shows the correlation between *XIST* expression and neXi gene expression levels in KS spermatogonia stem cells (SSCs). (F–I) Violin plot shows the percentage of total X chromosome (F), total Y chromosome (G), neXi (H), and eXi (I) gene expression levels in each germ cell subpopulation of OA and KS cells. (J) Volcano plot shows the correlation between neXi genes and each gene expression level in all KS SSCs. The X-axis represents the r value of the correlation coefficient. The Y-axis represents the P value. The sizes of the points represent the fold change of the gene expression level in KS cells compared with OA cells. (K) Regression analysis shows the correlation of the expression of the top six genes that are most strongly associated with neXi gene expression levels in KS SSCs. (L) Violin plot shows the expression of the top six genes that are most strongly associated with neXi gene expression levels in KS SSCs. (M) A violin plot shows the expression of the PRC1 and PRC2 complex in OA and KS germ cells.

## Discussion

KS is a common cause of male infertility and affects up to 3.1% of the infertile male population and approximately 10% of azoospermia patients(Lanfranco et al. 2004; Dabaja and Schlegel 2012). Testosterone replacement can improve the quality of life but has no positive effect on infertility, indicating that androgen deficiency may be not the main cause of spermatogenesis failure in KS patients. However, approximately 75% of adult males with KS are misdiagnosed because of normal fertility and a mild disease phenotype. In addition, most patients with azoospermia caused by KS (72%) have a possibility of sperm retrieval via microdissection and testicular sperm extraction(Dabaja and Schlegel 2012). These findings indicate that the dosage effect of the X chromosome may affect the progression and prognosis of KS. The dosage effect of the X chromosome in KS at least includes three aspects: 1) high-grade chromosome aneuploidies (48, XXXY; 48, XXYY; 49, XXXXY) or 46, XY/47, XXY mosaicism; 2) differences in sensitivities to X chromosome genes; and 3) differences in the ability to silence the extra X chromosome. Among these factors, the karyotype is arguably the most critical. It is reported that only 46, XY SSCs can enter into meiosis(Bergère et al. 2002). In this study, all three of our patients underwent successful sperm extraction, suggesting possible germ cell mosaicism. However, the DNA-FISH results showed that nearly all somatic cells in the seminiferous tubules, as well as those in vitreous degeneration lesions, had a karyotype of 47, XXY. With traditional RNA sequencing (RNA-seq), it is difficult to clearly describe the heterogeneous phenotype and explore the potential mechanism within KS testicular cells. Here, we profiled over 35,000 testicular single-cell transcriptomes from OA and KS patients and observed the heterogeneity of susceptibility to an extra X chromosome and the ability to counter X gene overload within KS testicular cells. Except in cells undergoing meiosis, XCI triggered by *XIST* is the most important regulatory mechanism of X chromosome silencing in XX cells. *XIST* RNA is transcribed in the XIC and then spreads to spatially close sites and eventually to the entire X chromosome(Galupa and Heard 2015). During the silencing maintenance phase, *XIST* RNA recruits silencing complexes, such as PRC1, PRC2, and Jarid2 to lock the inactive state of the post-XCI chromosome(Galupa and Heard 2015; Vidal and Starowicz 2017). Previous studies have mainly focused on females, and the researchers found and identified approximately 15% of X chromosome genes that may escape from XCI(Berletch et al. 2011). Our study obtained a similar result that most eXi genes were upregulated in KS somatic cells; however, the percentage of neXi gene expression in KS Sertoli cells was also increased over 1.5-fold. Furthermore, *XIST* showed low expression levels in KS Sertoli cells; however, in the few KS Sertoli cells that expressed *XIST*, a significant negative correlation was found between the *XIST* expression level and the percentage of neXi gene expression, indicating that the decreased XCI ability in KS Sertoli cells was caused by the quantity of *XIST* expression but not the dysfunction of the *XIST* mechanism. In addition, premeiotic germ cells also expressed low *XIST* levels similar to Sertoli cells, but they also expressed high levels of PRC1/2 complexes. In these cells, *XIST* expression was not negatively correlated with the percentage of neXi gene expression. Although the recruitment of PRC2 to the X chromosome is directly or indirectly depend on *XIST*, a significant spatial separation of PRC2 proteins and *XIST* was observed(Cerase et al. 2014). Furthermore, no PRC2 factors were found in a recent study purifying *XIST* interacting partners(Chu et al. 2015). Therefore, we speculated that premeiotic germ cells were in an *XIST*-independent XCI maintenance phase. We found that XCI is completely dominated by a different mechanism in meiotic cells, MSUC plays a critical role in both OA and KS cells.

*XIST* is tightly controlled by a series of pluripotency factors, trans-acting factors, and cis-regulators in an X dosage-sensitive manner. For example, YY1, which has been implicated as a trans-acting activator of *XIST* expression and helps to tether *XIST* RNA to the X chromosome during XCI (Jeon and Lee 2011), was widely expressed in different types of testicular cells but not in Sertoli cells. JPX, which enhances *XIST* expression in cis (Tian et al. 2010), was expressed in KS somatic cells but not in KS Sertoli cells and normal somatic cells. In addition, we found that most *XIST* regulators showed a “low expression in Sertoli cells” pattern, indicating the presence of one or more common upstream regulatory signaling disorders in KS Sertoli cells. Overexpression of *XIST* in Sertoli cells is a potential treatment for KS patients; however, *XIST* RNA is too long (19k bp) to embed in a vector. Therefore, a more feasible treatment may be to target upstream signaling of *XIST*. In addition, although KS Sertoli cells lack *XIST*, the neXi genes were not increased by two-fold in these cells, indicating the presence of other potential X-silencing mechanisms such as the PRC1/2 complex.

The pathological changes of KS Sertoli cells may be caused by abnormal estrogen receptor signaling and immune response as the pathway analysis. Furthermore, according to MGI database (http://www.informatics.jax.org/), we found 14 neXi_DEGs between OA and KS Sertoli cells have fertility associated phenotypes in mouse model (Table S2). For example, TIMP-1 function as a coregulator of basal testicular steroidogenesis, its mutant mice have higher testis weights than age-matched wild-type males(Nothnick et al. 1998); UTX (H3K27me3 demethylase) may cause down-regulation of lactate and cholesterol metabolism so that affects the nursing functions of XX/Sry Sertoli cells(Shishido et al. 2017); HPRT1 plays a central role in the generation of purine nucleotides through the purine salvage pathway, its mutant male mice at 34-38 weeks displayed complete testicular atrophy(Ordway et al. 1997). The abnormal high expression of these neXi_DEGs may directly damage Sertoli cells alone or coordinately, and caused spermatogenesis failure.

Although we performed a transcriptional analysis to uncover the pathological changes and potential mechanisms of the KS spermatogenic microenvironment, additional evidence is still needed, such as immunohistochemistry and RNA-FISH in testis tissue sections. Some candidate regulators also need to be identified by functional experiments. In addition, some important regulators of *XIST* such as its antisense version, *TS/X*, were not detected in this study because its RNA lack 3’ poly-A signaling to be captured and underwent reverse transcription. We observed abnormal hormonal and immune responses in KS Sertoli cells and predicted that some small molecule drugs could potentially have therapeutic effects, such as ST 1962, sirolimus, and CD437. In the future, the spermatogenic microenvironment in KS patients may be improved through endocrine regulation and suppression of abnormal immune responses so that genetic safety issues caused by gene editing can be avoided.

In general, based on the single-cell transcriptome data from normal adults and KS patients, we propose a novel mechanism in which the weak XCI ability of Sertoli cells causes their high susceptibility to the extra X chromosome, and this increased damage to Sertoli cells leads to atrophy of seminiferous tubules and failure of spermatogenesis.

---

**Figure S1.**
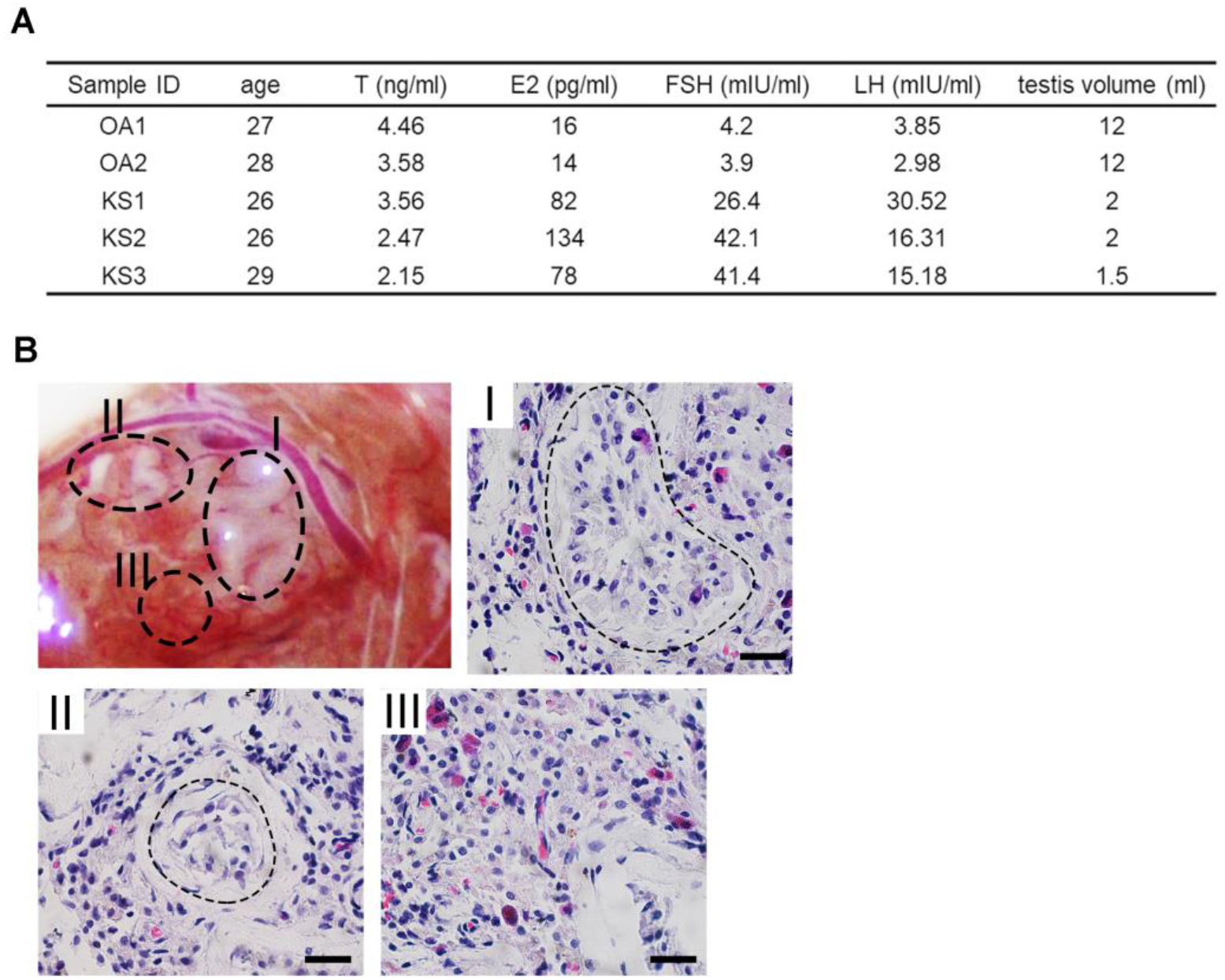
Clinical information and Heterogeneous testicular atrophy in Klinefelter syndrome. (A) The clinical information of 2 OA and 3 KS patients. (B) Photograph of a typical KS testis during microdissection and testicular sperm extraction revealing heterogeneous atrophy in different regions. Hematoxylin and eosin staining shows the histological changes in each region. Region I, larger and opaque seminiferous tubules with normal spermatogenesis; Region II, thin and transparent seminiferous tubules that only contain pathological Sertoli cells; Region III, severely atrophic seminiferous tubules with almost all Sertoli cells and lacking germ cells. The scale bar represents 40 μm.

**Figure S2.**
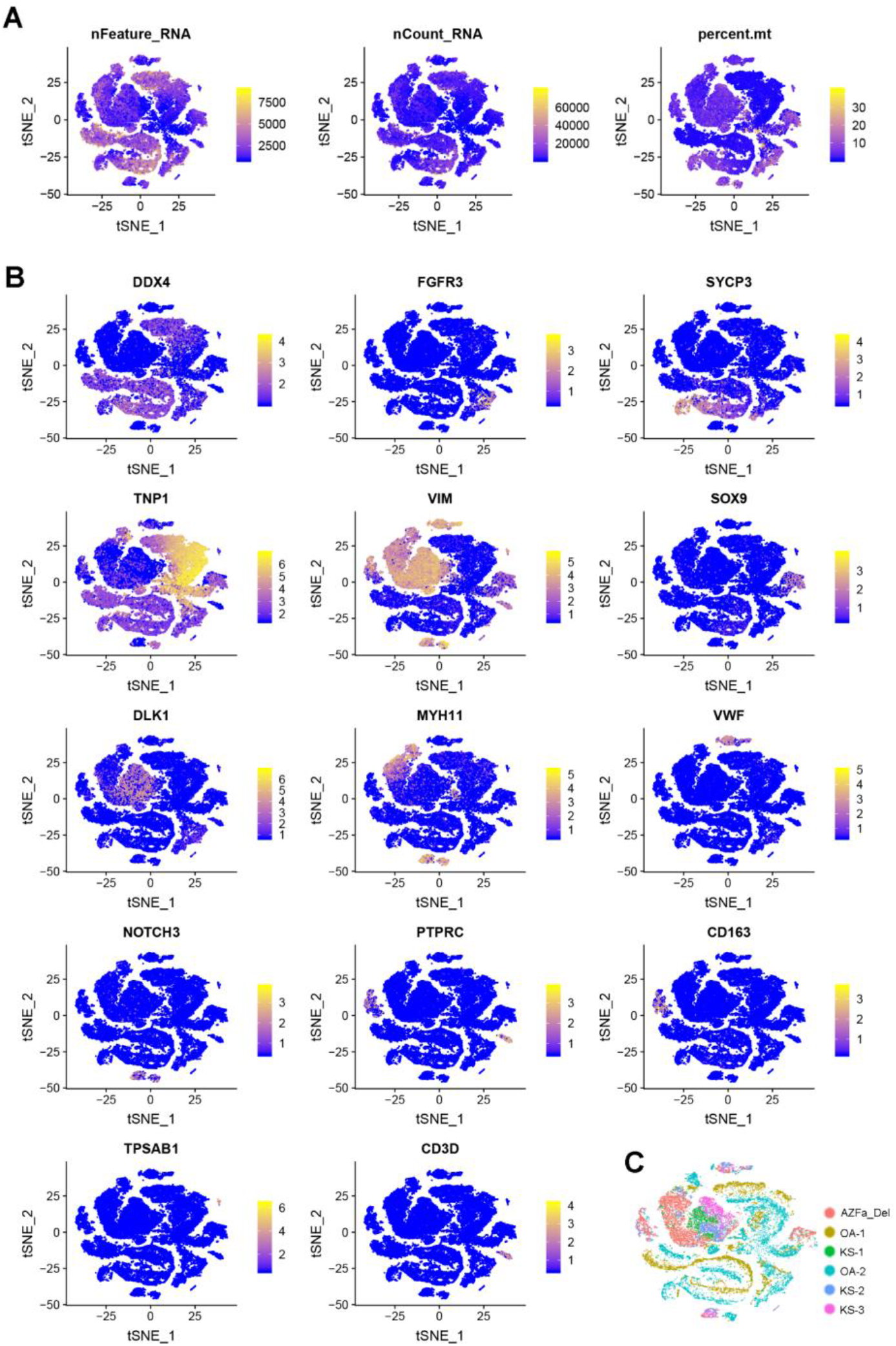
Single-cell RNA sequencing quality information and known marker expression in each cluster. (A) The single-cell RNA sequencing quality information includes the total gene expression counts, unique molecular identifier counts, and the percent of mitochondrial gene expression projected on the T-distributed stochastic neighbor embedding (tSNE) plot. (B) Expression patterns of selected markers projected on the tSNE plot.

**Figure S3.**
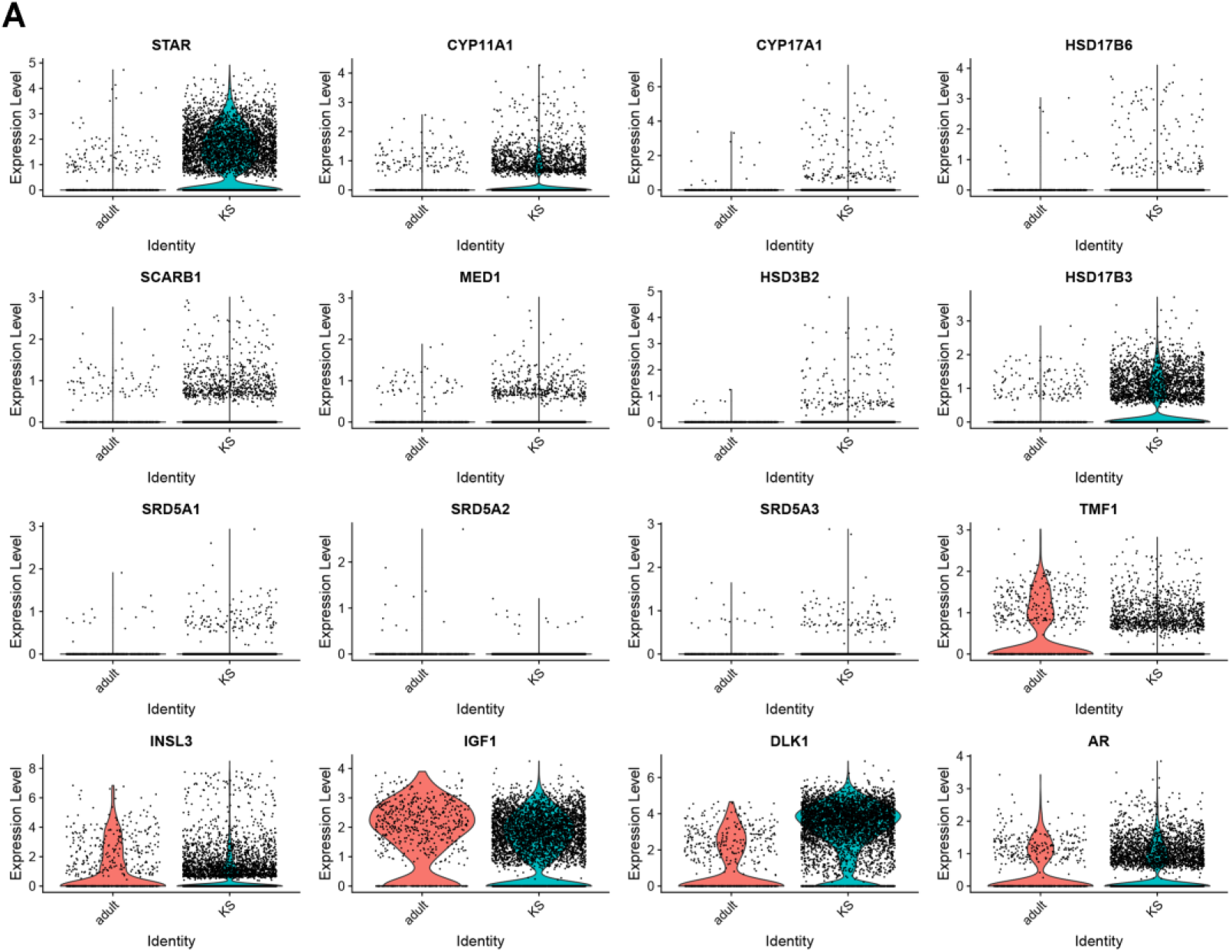
Change in the androgen synthesis pathway in Klinefelter syndrome Leydig cells. (A) Violin plot shows the expression levels of androgen synthesis pathway-related genes in Klinefelter syndrome Leydig cells.

## Methods

### Contact information for reagents and resource sharing

The RNA-seq matrix data can be obtained from the GEO database (GSE149512). Requests for further information, resources, and reagents should be directed to and will be fulfilled by the lead contact, Zheng Li.

### Experimental model and subjects

The experiments performed in this study were approved by the Ethics Committee of Shanghai General Hospital (License No. 2016KY196). All participants signed consent forms after being fully informed of the goal and characteristics of our study. Fresh testicular tissues were obtained from abandoned tissues after testicular sperm extraction operations performed for the following indications: OA (*n*=2), KS (*n*=3) and YqAZFa microdeletion (*n*=1). The karyotypes, genotypes, sex hormone levels, and morphology of seminiferous tubules of OA donors were normal. The karyotypes of three KS samples were XXY without mosaic. The YqAZFa microdeletion sample was used as a negative control to exclude germ cell interference. This patient was diagnosed by qPCR examination before hospitalization, which showed that in this sample, sY84 and sY86 were completely deleted. Among all donors, other abnormal genotypes related to spermatogenic disorders were excluded by whole-exome sequencing.

### Histological examination

Fresh testicular tissues from donors were fixed in 4% paraformaldehyde for 12–24 hours at 4°C, embedded in paraffin, and sectioned. Before staining, tissue sections were dewaxed in xylene, rehydrated using a gradient series of ethanol solutions, and washed in distilled water. Then, the sections were stained with PAS/hematoxylin and dehydrated using increasing concentrations of ethanol and xylene. Sections were allowed to dry before applying neutral resin to the coverslips. The staining images were captured with a Nikon Eclipse Ti-S fluorescence microscope.

### DNA-FISH of KS Sertoli cells

KS Sertoli cells were isolated by sequential two-step enzyme digestion as described in our previous study(Yang et al. 2019). Briefly, testicular tissues were enzymatically digested with 4 mg/ml collagenase type IV for 5 min to remove most interstitial cells (such as Leydig cells and PTM cells); then, the tissues were digested with 4 mg/ml collagenase type IV, 2.5 mg/ml hyaluronidase, and 1 mg/ml trypsin at 37°C for 20 min to isolate the remaining tissue as single cells. After adherent growth for 24 hours, these cells were re-digested with 4 mg/ml collagenase type IV for 1 min to remove germ cells. The remaining adherent Sertoli cells were digested with trypsin and fixed with 4% paraformaldehyde. DNA-FISH was performed according to the guidelines of the CEP X SpectrumOrange/Y SpectrumGreen DNA Probe Kit (Abbott, 07J20-050 and 07J20-050). In brief, after denaturation of the specimen DNA at 73°C for 5 min, 10 μl of the probe solution was incubated with the slide at 42°C for 30 min. After washing with SSC buffer (sodium chloride and sodium citrate), nuclei were labeled with DAPI by incubating tissue sections for 15 min. Images were captured with an OLYMPUS IX83 confocal microscope.

### Mapping, sample quality control, and integration of single-cell RNA-seq data analysis

Single-cell isolation and RNA-seq library preparation were described in our previous study(Zhao et al. 2020). The raw data are labeled as GSE149512. After mapping, a Seurat object was created using the gene expression matrix via the Seurat package in R. Cells were further filtered according to the following threshold parameters: total number of expressed genes, 500 to 9000; total UMI count, between - ∞ and 35,000; and proportion of mitochondrial genes expressed, <40%. Normalization was performed according to the software package manual (https://satijalab.org/seurat/v3.1/pbmc3k_tutorial.html). Batch correction was performed using the IntegrateData function in the Seurat package.

### Cell identification and clustering analysis

After batch correction, the merged Seurat objects were scaled and analyzed by principal component analysis. Then, the first 20 principal components (PCs) were used to construct a KNN graph and refine the edge weights between any two cells. Based on all of the local cell neighborhoods, the FindClusters function with a resolution parameter of 0.1 was used to cluster the cells. In total, 17 clusters were identified, and these clusters were renamed according to accepted marker genes. The first 20 PCs were also used to perform non-linear dimensional reduction by tSNE, and the dimension reduction plots were produced as output (Figure 1B, 1C, and S2).

### Similarity and correlation of different types of KS and OA or AZFa_Del testicular cells

The expression of each gene was treated as an independent variable. Similarities were represented with the Jaccard index, which was calculated with the vegan 2.5-6 R package. The result is shown as 1-Jaccard distance. The correlations were calculated with the function cor(method=spearman) in the stats 3.6.0 R package.

### Identification of DEGs

The Seurat FindAllMarkers function (test.use=“wilcox”) is based on the normalized UMI count and was used to identify unique cluster-specific marker genes. Only the genes that were detected in at least 10% of the cells were tested, and the average log2 (fold change) threshold was set as 0.25, 0.5, or 1 as needed.

### Upstream regulators, canonical pathways, and bio-function analysis

IPA was used to analyze the upstream regulators, canonical pathways, and bio-function differences between KS and OA Sertoli cells. A total of 1333 DEGs with a change of over 0.25 log fold change and with a P value less than 0.05 were input into IPA. IPA terms with a P value less than 0.05 and a z score higher/less than 2/-2 were considered significant.

### Prediction and analysis of candidate transcriptional regulators of *XIST*

The first 3000 bases at the transcription initiation site of *XIST* were considered the regulatory binding region, and 23 potential transcription regulators were predicted through the PROMO database. Then, the binding sites of these transcription factors were found and scored using the JASPAR database.

### Transcription factor network construction

A total of 1469 human transcription factors in the AnimalTFDB data set and a gene expression matrix of OA and KS Sertoli cells were used as input for the ARACNe-AP software. The results of the ARACNe-AP analysis were input into the ssmarina (version 1.01) R package, which further calculated the marina objects containing the normalized enrichment scores, P values, and a specific set of regulators.

## Acknowledgment

This work was support by grants from National Key R&D Plan of China (2017YFC1002003 and 2018YFA0107702); National Natural Science Foundation of China (81671512, 81701524 and 81701428); the ShanghaiTech University start-up fund of Zhi Zhou; Doctorial innovation fund of Shanghai Jiao Tong University School of Medicine (BXJ201940); Project funded by China Postdoctoral Science Foundation (2019M661524). We wish to thank the (1) Shanghai OE Biotech CO. LTD, (2) Sinotech Genomics CO. LTD. Shanghai and (3) Novogene Bioinformatics Institute, Beijing, China for their support of RNA-seq library preparation. We thank Lisa Kreiner, PhD, from Liwen Bianji, Edanz Editing China (www.liwenbianji.cn/ac), for editing the English text of a draft of this manuscript.

